# Characterization of spike processing and entry mechanisms of seasonal human coronaviruses NL63, 229E and HKU1

**DOI:** 10.1101/2024.04.12.589332

**Authors:** Sabari Nath Neerukonda, Russell Vassell, Sabrina Lusvarghi, Shufeng Liu, Adovi Akue, Mark KuKuruga, Tony T. Wang, Carol D Weiss, Wei Wang

**Affiliations:** Office of Vaccine Research and Review, Center for Biologics Evaluation and Research, US Food and Drug Administration, Silver Spring, Maryland

**Keywords:** seasonal coronavirus, neutralization, pseudovirus, virus entry, transmembrane serine protease 2

## Abstract

Although much has been learned about the entry mechanism of severe acute respiratory syndrome coronavirus-2 (SARS-CoV-2), the details of entry mechanisms of seasonal human coronaviruses (HCoVs) remain less well understood. In the present study, we established that 293T cell lines that stably express angiotensin converting enzyme (ACE2), aminopeptidase N (APN), or transmembrane serine protease 2 (TMPRSS2) support high level transduction of lentiviral pseudoviruses bearing spike proteins of seasonal HCoVs, HCoV-NL63, -229E, or -HKU1, respectively. Our results showed that entry of HCoV-NL63, -229E and -HKU1 pseudoviruses is sensitive to endosomal acidification inhibitors (chloroquine and NH_4_Cl), indicating virus entry via the endocytosis route. Although HCoV-HKU1 pseudovirus infection requires TMPRSS2 expression on cell surface, endocytosis-mediated HCoV-HKU1 entry requires the serine protease domain but not the serine protease activity of TMPRSS2. We also show that amino acids in the predicted S1/S2 junctions of spike proteins of HCoV-NL63, and - 229E are essential for optimal entry but non-essential for spike-mediated entry of HCoV-HKU1. Our findings provide insights into entry mechanism of seasonal HCoVs that may support the development of novel treatment strategies.

**Importance:** Details of the entry mechanisms of seasonal human coronaviruses (HCoVs) remain to be fully explored. To investigate the entry of HCoV-NL63, -229E and -HKU1 CoVs, we employed 293T cells that stably express angiotensin converting enzyme (ACE2) aminopeptidase N (APN), or transmembrane serine protease 2 (TMPRSS2) to study entry mechanisms of pseudoviruses bearing spike proteins of HCoV-NL63, -229E and - HKU1 respectively. Our results provide new insights into the predicted S1/S2 subunit junctions, cellular receptor, and protease requirements for seasonal HCoV pseudovirus entry via endocytic route and may support the development of novel treatment strategies.

## Introduction

The coronavirus disease 2019 (COVID-19) global pandemic, caused by Severe Acute Respiratory Syndrome Coronavirus 2 (SARS-CoV-2), has rekindled interest in spike protein functions of seasonal human coronaviruses (HCoVs). The *Coronavirinae* subfamily has four seasonal HCoVs that are frequently associated with mild upper respiratory disease in immunocompetent individuals but may rarely be associated with severe disease in comorbid individuals with underlying disease conditions or those that are immunocompromised (1). Seasonal HCoVs include *alphacoronavirus* (HCoV-NL63 and -229E) and *betacoronavirus* (HCoV-HKU1 and -OC43) genera members.

Seasonal HCoVs, also known as “common cold” coronaviruses, infect individuals within the first two decades of life with reinfections occurring throughout life. Although seasonal HCoVs exhibit sustained endemic and widespread transmission, they tend to exhibit biennial pattern of seasonality with at least one *alphacoronavirus* and one *betacoronavirus* cocirculating each season (1). Biennial pattern of seasonality is supported by serologic and experimental human studies indicating short-lived immunity that lasts at least a year, prior to reinfection due to waning immunity (2, 3) or genetic drift (4, 5). Several key questions regarding seasonal HCoV immunity are being addressed, including the role of pre-existing seasonal HCoV antibodies (Abs) in protection against COVID-19, immune escape of seasonal HCoVs (4–6), potency of broadly neutralizing Abs (7, 8), and Ab durability following natural infection or vaccination (2). While Ab binding may not always correlate with protection against clinical disease, neutralizing Abs (nAbs) impeding receptor binding and/or fusion steps of entry often correlate with protection against clinical disease. Therefore, a fuller understanding of spike-mediated entry mechanisms will facilitate the development of therapeutic nAbs and nanobody entry inhibitors (4–6, 9).

All coronavirus spike glycoproteins have similar domain organization, with a furin processing site at the S1 and S2 subunit junction, followed by a S2’ processing site that gets cleaved by cellular proteases during entry. The S1 subunit consists of an N-terminal domain that facilitates cellular attachment and a C-terminal domain harboring receptor-binding domain (RBD). The S2 subunit consists of domains necessary for fusion including fusion peptide (FP), heptad repeats (HR) 1 and 2 and stem helices followed by a transmembrane domain and cytoplasmic tail.

Although a few HIV lentivirus-based pseudotyping systems have been described for seasonal HCoV spikes (7, 10), entry mechanisms of seasonal HCoVs pseudoviruses have not been fully explored. Here, we established optimal infection conditions in 293T cells that stably express functional receptors for seasonal HCoV spike-mediated entry and investigated the functional requirement of cellular protease and furin cleavage sites in the S1/S2 junction for spike-mediated entry and the routes of entry followed by seasonal HCoVs.

## Results

### Functional requirement of furin processing site for entry of HCoV-NL63, 229E- and HKU1 pseudoviruses in target cells

To investigate spike-mediated entry of seasonal HCoVs, we established 293T cells that stably expressed ACE2 and APN or TMPRSS2 and used 293T-ACE2 cells established by Crawford *et al* (11). As expected, pseudoviruses bearing full-length spikes of HCoV-NL63, -229E, and -HKU1 underwent high levels of infection in 293T cells that stably express ACE2, APN, and TMPRSS2, respectively, (Supplementary Figures 1A-C), whereas SARS-CoV-2 control displayed high level infectivity in 293T cells that express both ACE2 and TMPRSS2 (Supplementary Figure 1D) (12). To confirm the specificity of entry due to functional interaction between S1 subunit and cellular receptor, we inhibited entry step using rabbit sera raised against S1 subunit of each seasonal HCoV spike. Rabbit sera raised against seasonal HCoV spike components neutralized homologous seasonal HCoV, confirming specificity of infection in the respective target cells (Supplementary Figure 2).

We investigated the role of furin processing sites in HCoV-NL63, -229E and - HKU1 entry by modifying predicted furin cleavage sites (FCS) in S1/S2 junctions. The spike proteins of HCoV-NL63, -229E and -HKU1 were predicted to have dibasic (SLIPVRPR|SS; ‘|’ represents furin proteolytic cleavage junction), monobasic (SIIAVQPR|NVS), and multibasic (SSSRRKRR|SIS) FCS in S1/S2 junction respectively (13, 14). We engineered deletions into HCoV-NL63 spike expression plasmid to express spike protein with either a monobasic FCS (mFCS; SLIPVR|SS) or deleted FCS (ΔFCS; (SLIPV|SS). Compared to pseudoviruses bearing wild type (WT) spike, pseudoviruses bearing ΔFCS spike displayed a 15-fold reduction in infectivity whereas pseudoviruses bearing mFCS spike lacked infectivity (Figure 1A). Deletion of residues in FCS did not abolish furin processing of spike into S1 subunit in pseudoviruses bearing mFCS and ΔFCS spikes, but modestly increased the levels of full-length spike compared to pseudoviruses bearing WT spike (Figure 1B). These results demonstrate that FCS modifications are deleterious to HCoV-NL63 spike-mediated infectivity. Furthermore, HCoV-NL63 spike underwent efficient cleavage despite P4-P1 positions of the predicted FCS (VRPR) not conforming to standard FCS sequence pattern (R-X-[K/R]-R), suggesting an alternative FCS in spike protein might allow efficient processing into S1/S2 subunits (15, 16).

**Figure 1.**
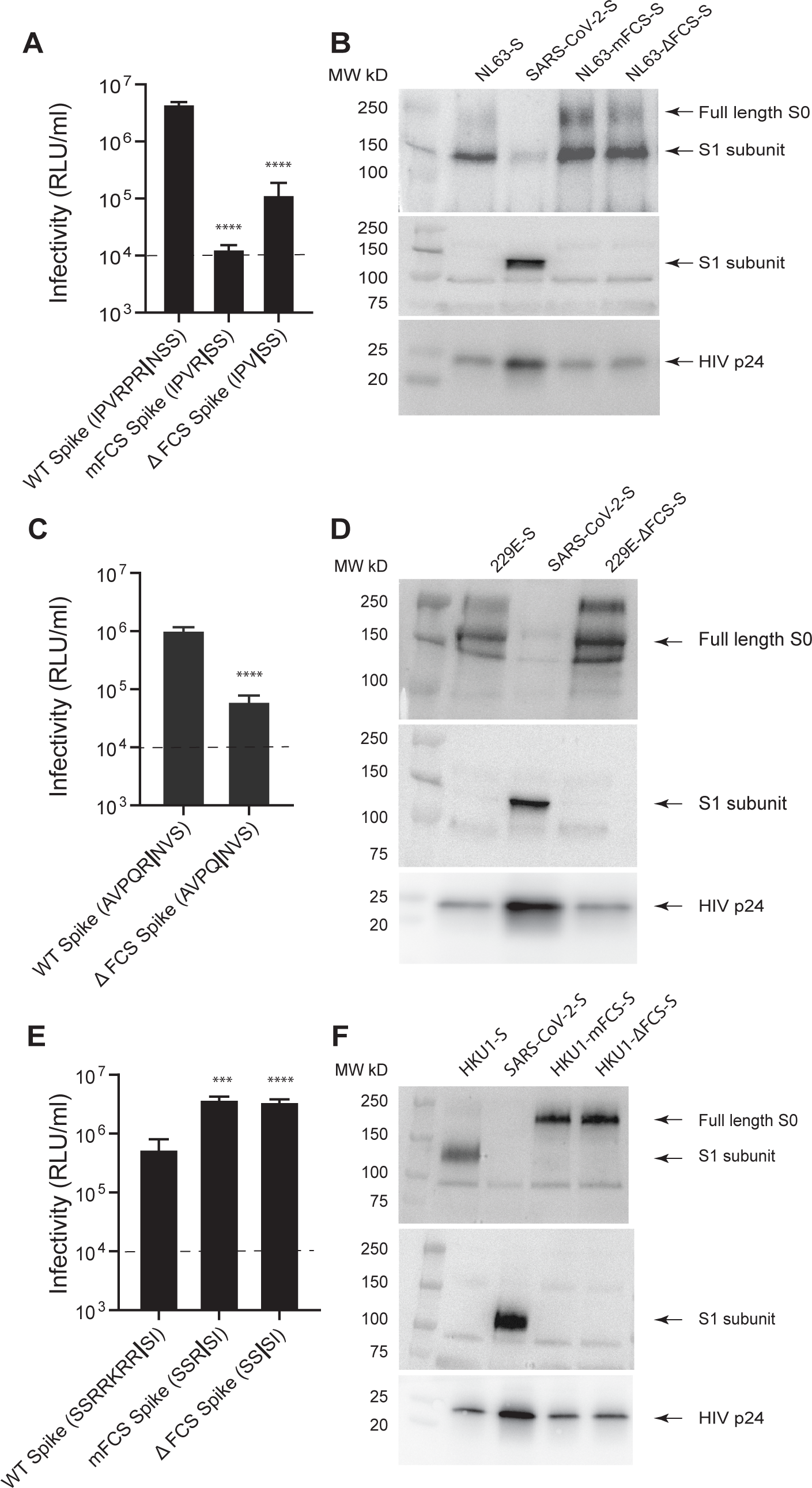
Functional impact of predicted furin processing sites on pseudovirus infectivity and spike expression. Infectivity of pseudoviruses bearing wild type (WT), monobasic furin cleavage site (mFCS) and deleted furin cleavage site (ΔFCS) spike proteins of HCoV-NL63 (A), -229E (C), and -HKU1 (E) in 293T-ACE2, 293T-ACE2-APN, and 293T-TMPRSS2. Dotted line indicates background level (1 × 10^4^ RLU/ml). Data are shown as means and standard deviations from three independent experiments. The tests for two-group comparison (mutant spike vs WT spike) were analyzed using GraphPad Prism software. Asterisk* denotes significance: ****: *p ≤ 0.0001*. Western blotting of HCoV-NL63 (B), -229E (D), -HKU1 (F) and SARS-CoV-2 pseudoviruses (B, D and F) bearing indicated spike proteins. HCoV-NL63 (B), -229E (D), -HKU1 (F) and SARS-CoV-2 spike proteins were probed using anti-NL63-S1 mouse mAb (9D3F3), anti-229E-S1 mouse mAb (24C9F7), anti-HKU1-S1 rabbit polyclonal sera and rabbit pAb (Sino Biological, 40589-T62) respectively, Full length S0 and furin processed S1 subunits were indicated using arrows. HIV p24 probed using mouse hybridoma (183-H12-5) is shown for pseudovirus control.

We next engineered one amino acid deletion in HCoV-229E spike to generate ΔFCS spike (SIIAVQP|NVS). Compared to pseudoviruses bearing WT HCoV-229E spike, pseudoviruses bearing ΔFCS spike has 17-fold lower infectivity, suggesting FCS modification described here is deleterious for spike-mediated entry and infectivity (Figure 1C). Both HCoV-229E WT and ΔFCS spike bearing pseudoviruses expressed full length spike protein with a predicted size of ∼170kD, and therefore remained largely uncleaved by the furin (Figure 1D). These findings support nonconformity of predicted FCS (AVQP) of HCoV-229E spike with the standard FCS sequence pattern (R-X-[K/R]-R) and therefore lack of furin cleavage (15, 16).

We next engineered deletions in the multibasic FCS (RRKRR|SIS) of HCoV-HKU1 spike to generate mFCS (SSSR|SIS) and ΔFCS (SSS|SIS) spikes. Unexpectedly, pseudoviruses bearing mFCS and ΔFCS spikes had 7-fold and 6.4-fold higher infectivity, respectively, than WT HCoV-HKU1 spike (Figure 1E). HCoV-HKU1 pseudoviruses bearing WT spike protein displayed efficient processing into S1 subunit (∼125kD), whereas pseudoviruses bearing mFCS and ΔFCS spike proteins failed to undergo furin cleavage as shown by the presence of bands corresponding to the size of full-length spike protein (∼200kD) (Figure 1F). The P4-P1 positions in the FCS of HCoV-HKU1 spike (RKRR) conforms to the standard FCS sequence pattern (R-X-[K/R]-R), and therefore underwent efficient furin cleavage. Deletion of FCS abolished furin cleavage of HCoV-HKU1 spike (15, 16).

### HCoV-NL63, -229E and -HKU1 pseudoviruses follow endocytic route of entry

We next employed chemical inhibitors that either inhibit endosomal acidification (chloroquine, Ammonium chloride or NH_4_Cl) or TMPRSS2 catalytic activity (Camostat mesylate) to determine the route of entry and the requirement of TMPRSS2 serine protease during entry respectively. In 293T-ACE2 cells lacking TMPRSS2, HCoV-NL63 pseudovirus entry is sensitive to chloroquine inhibition (IC_50_, 7.05 μM) and therefore, entered via endosomal route of entry as previously described for SARS-CoV-2 D614G pseudoviruses (IC_50_, 3.84 μM) and shown here as a positive control (Figure 2A). SARS-CoV-2 pseudovirus entry is sensitive to camostat mesylate in 293T-ACE-TMPRSS2 cells (IC_50_, 0.94 μM) but not in 293T-ACE2 cells as previously described (Figures 2B) (12). By contrast, HCoV-NL63 pseudovirus entry remained sensitive to chloroquine inhibition (IC_50_, 5.66 μM) in 293T-ACE2-TMPRSS2 cells (Figures 2A) and showed lack of sensitivity to camostat mesylate (Figures 2B), indicating that HCoV-NL63 spike-mediated entry follows endosomal route and is independent of TMPRSS2 protease activity. These findings concur with earlier infection results (Supplementary Figure 1A) where no enhancement of HCoV-NL63 pseudovirus infection was observed in 293T-ACE2-TMPRSS2 cells compared to 293T-ACE2 cells, further confirming that TMPRSS2 is dispensable for HCoV-NL63 spike-mediated entry.

**Figure 2.**
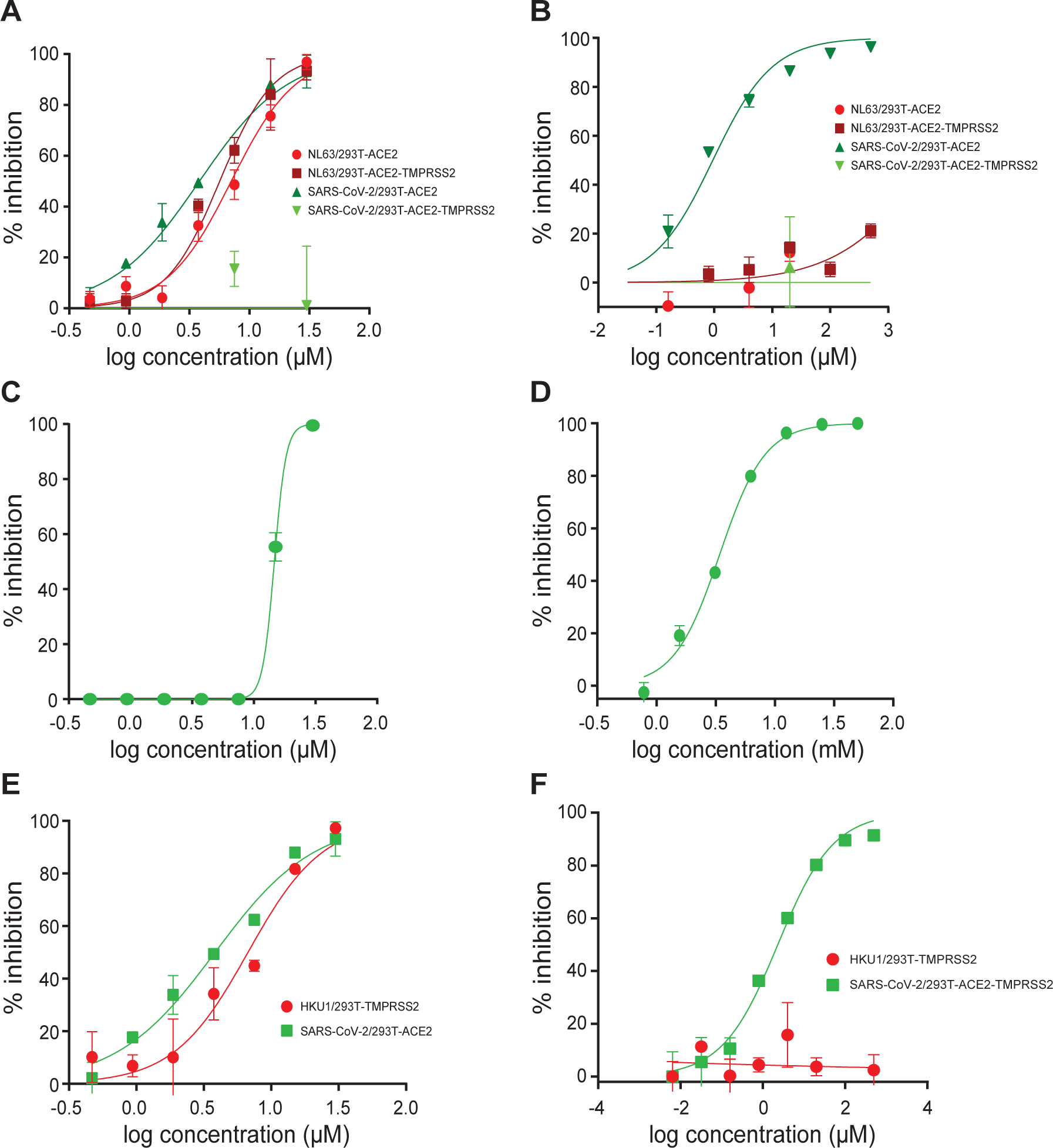
Endosomal entry of HCoV-NL63, -229E, and -HKU1 pseudoviruses. (A) Chloroquine sensitivity of HCoV-NL63 and SARS-CoV-2 pseudoviruses in 293T-ACE2 and 293T-ACE2-TMPRSS2 cells. (B) Camostat mesylate sensitivity of HCoV-NL63 and SARS-CoV-2 pseudoviruses in 293T-ACE2 and 293T-ACE2-TMPRSS2 cells. (C) Chloroquine sensitivity of HCoV-229E pseudoviruses in 293T-ACE2-APN cells. (D) Ammonium chloride sensitivity of HCoV-229E pseudoviruses in 293T-ACE2-APN cells. (E) Chloroquine sensitivity of HCoV-HKU1 and SARS-CoV-2 pseudoviruses in 293T-TMPRSS2 and 293T-ACE2 cells. (F) Camostat mesylate sensitivity of HCoV-HKU1 and SARS-CoV-2 pseudoviruses in 293T-TMPRSS2 and 293T-ACE2-TMPRSS2 cells respectively. For all experiments, cells were pretreated with indicated inhibitor for 2 h prior to pseudovirus infection in the presence of inhibitor for a period of 48 h. X-axis indicates log concentration of inhibitor. Y axis indicates percentage inhibition compared to pseudovirus infection without inhibitor treatment. Results shown are representative of three independent experiments.

We next investigated entry route of HCoV-229E pseudoviruses in 293T-ACE2-APN cells by examining their sensitivity to chloroquine, NH_4_Cl and camostat mesylate inhibition. HCoV-229E entry is sensitive to chloroquine (IC_50_, 14.65 μM) (Figures 2C) and NH_4_Cl (IC_50_, 3.41 mM) (Figures 2D), indicating entry via endosomal route. These results along with infectivity results (Supplementary figure 1B) described earlier indicate that TMPRSS2 expression is not required for HCoV-229E pseudovirus entry, since 293T-ACE2-APN cells support HCoV-229E pseudovirus entry without TMPRSS2 coexpression.

We next determined the entry route of HCoV-HKU1 pseudoviruses in 293T-TMPRSS2 cells by examining their sensitivity to chloroquine, NH_4_Cl and camostat mesylate inhibition. Surprisingly, although HCoV-HKU1 pseudovirus infection required TMPRSS2 expression (Supplementary Figure 1C) (10), HCoV-HKU1 entry remained sensitive to chloroquine (IC_50_, 7.34 μM) (Figures 2E) and NH_4_Cl (IC_50_, 1.73 mM) (Supplemental Figure 3), but not camostat mesylate (Figures 2F). These results indicate that HCoV-HKU1 entry occurred via endosomal route. As a control, SARS-CoV-2 entry is sensitive to chloroquine (Figure 2E) or NH_4_Cl (IC_50_, 1.5 mM) (Supplemental Figure 3) inhibition in 293T-ACE2 cells and camostat mesylate inhibition (Figure 2F) in 293T-ACE2-TMPRSS2 cells as previously described (12). The lack of sensitivity to camostat mesylate inhibition in HCoV-HKU1 entry highlights that TMPRSS2 facilitates HCoV-HKU1 entry via a mechanism that is independent of its serine protease activity. These findings led us to investigate whether HCoV-HKU1 spike-mediated entry is independent of TMPRSS2 serine protease activity or dependent on domains other than its serine protease (SP) domain.

### Role of non-catalytic TMPRSS2 in HCoV-HKU1 entry

TMPRSS2 is a type II transmembrane glycoprotein that belongs to the type 2 transmembrane serine protease (TTSP) family and is composed of an N-terminal intracellular domain, a single-pass transmembrane (TM) domain and an ectodomain comprising three subdomains: a low-density lipoprotein receptor type A (LDLRA) domain, a Class A Scavenger Receptor Cysteine-Rich (SRCR) domain and a C-terminal trypsin-like Serine protease (SP) domain with a canonical H296-D345-S441 catalytic triad (Figure 3A) (17, 18). TMPRSS2 is synthesized as a single chain zymogen (∼65kD) and is intracellularly autoactivated at R^292^-I^293^ junction to generate cleaved protease fragment (∼31kD) and two additional lighter bands of (∼45 to 48 kDa), possibly generated upon cleavages between the LDLRA and the SRCR domains (19).

**Figure 3.**
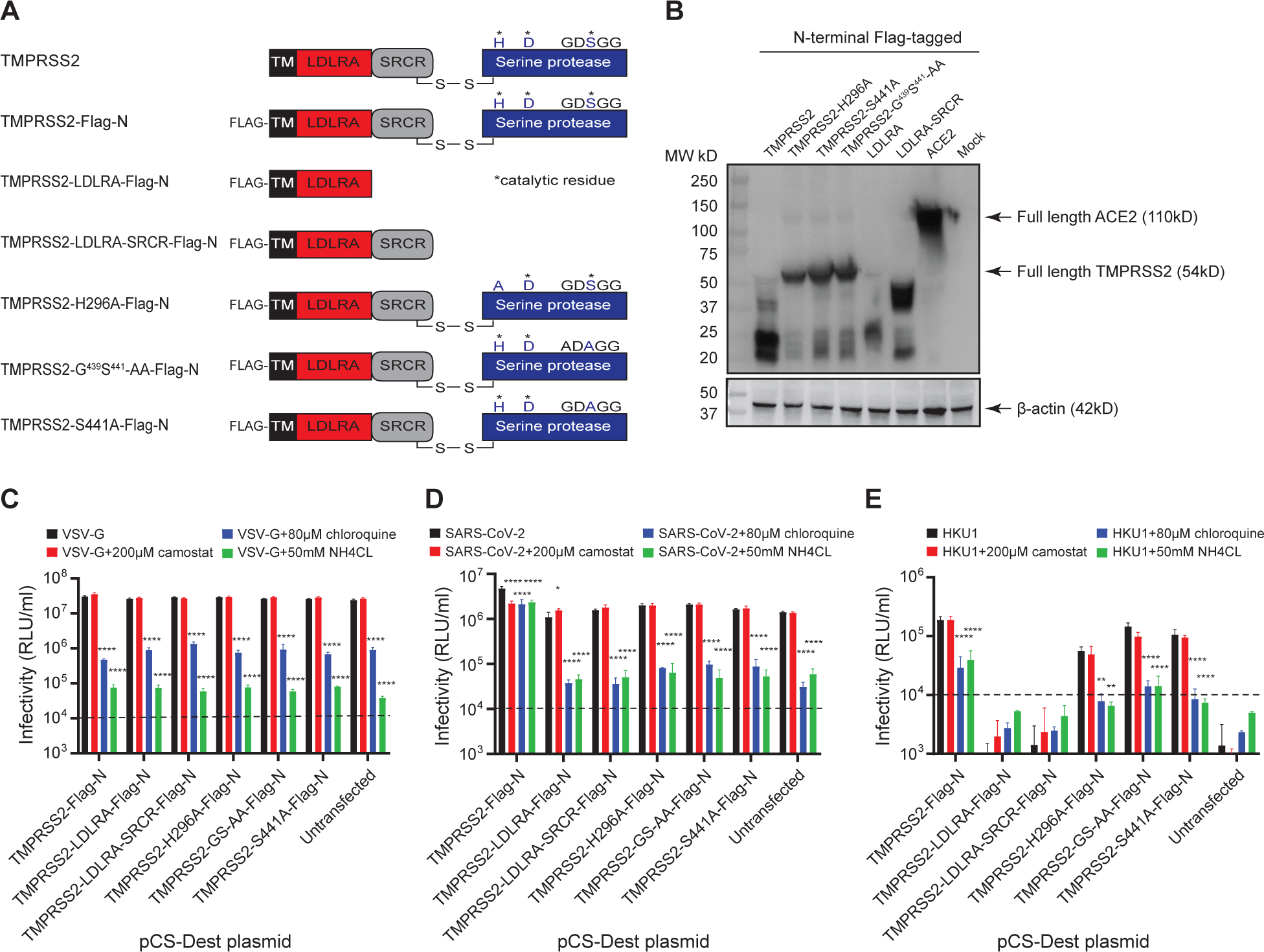
Functional domain organization and role of TMPRSS2 proteins in HCoV-HKU1 pseudovirus entry. (A) Cellular TMPRSS2 is a type II transmembrane protease comprising N-terminal transmembrane (TM) domain, Low Density Lipoprotein Receptor domain A (LDLRA), Class A Scavenger Receptor Cysteine-Rich (SRCR) domain and a C-terminal trypsin-like serine protease (SP) domain with a canonical H296-D345-S441 catalytic triad. TMPRSS2 with an N-terminal Flag tag, LDLRA domain of TMPRSS2 with an N-terminal Flag tag, LDLRA and SRCR domains of TMPRSS2 with an N-terminal Flag tag, and TMPRSS2 proteins with alanine substitutions in the indicated active site residues of catalytic triad are shown. (B) Western Blotting of Flag-tagged TMPRSS2 proteins in 293T cells transfected with indicated plasmids. Lane 1 is Marker, Lane 2 showed TMPRSS2 expressed as full-length form (∼54KD) and cleavage products (∼20-25KD, 37-40KD) due to autoactivation. Lane 3-5 showed TMPRSS2 proteins lacking protease catalytic acitivity expressed mainly as full-length forms (∼54KD). Lanes 6 and show LDLRA and LDLRA-SRCR domains expressed as 25KD and 37KD forms. Lane shows Flag-tagged ACE2 (∼110KD) as a control. (C) Infectivity of VSV pseudovirses in 293T cells transfected with indicated TMPRSS2 plasmids (black bars). Infectivity of VSV pseudoviruses in the presence of camostat, chloroquine and NH_4_Cl are indicated in red, blue, and green bars respectively (D) Infectivity of SARSCoV-2 pseudovirses in 293T-ACE2 cells transfected with indicated TMPRSS2 plasmids (black bars). Infectivity of SARS-CoV-2 pseudoviruses in the presence of camostat, chloroquine and NH_4_Cl are indicated in red, blue and green bars respectively (E) Infectivity of HCoV-HKU1 pseudovirses in 293T cells transfected with indicated TMPRSS2 plasmids (black bars). Infectivity of HCoV-HKU1 pseudoviruses in the presence of camostat, chloroquine and NH_4_Cl are indicated in red, blue and green bars respectively. The tests for two-group comparison (infectivity without inhibitor vs infectivity with inhibitor) were analyzed using GraphPad Prism software. Asterisk* denotes significance: *\*: p ≤ 0.05; **: p ≤ 0.01; ****: p ≤ 0.0001*.

To investigate if domains other than SP domain might support HCoV-HKU1 spike-mediated entry, we engineered stop codons in TMPRSS2-Flag-N plasmid within nucleotide sequences between LDLRA and SRCR (Y198 aa position), or SRCR and SP domains (R^292^I^293^ aa position) to allow expression of N-terminally Flag-tagged LDLRA or LDLRA-SRCR domains respectively (Figure 3A). To investigate if HCoV-HKU1 spike-mediated entry is independent of TMPRSS2 SP activity, we introduced single amino acid substitutions H296A and S441A in the catalytic triad of SP domain to abolish catalytic activity (Figure 3A) (19, 20). Structural basis of TMPRSS2 catalysis involves formation of an oxyanion hole by the backbone amine groups of G439 and S441 in the GDSGG motif to help activate and stabilize the carbonyl of the scissile bond (21). We therefore also generated a double amino acid mutant by introducing G439A and S441A substitutions (GS-AA) to abolish oxyanion hole formation and SP activity (Figure 3A).

Plasmids encoding LDLRA and LDLRA-SRCR domains, and TMPRSS2 proteins defective for SP catalytic activity (H296A, S441A and GS-AA) were transfected into 293T cells and confirmed for protein expression by Western Blotting (Figure 3B). In transfected 293T cells, full-length TMPRSS2 is expressed as ∼54kDa band which is autoactivated possibly at the R^292^-I^293^ junction to generate an N-terminal cleavage product of ∼24kDa and a cleaved protease fragment (∼31kDa) (17, 19). Since we probed for TMPRSS2 using an antibody directed against Flag tag at the N-terminus, full-length TMPRSS2 and several cleavage products were detected. A major N-terminal cleavage product of ∼24kDa possibly generated by autocleavage after LDLRA domain and additional lighter bands (∼37 to 40 kDa), possibly generated upon cleavages between SRCR and SP domains were detected (Figure 3B, Lane 2). Catalytically inactive TMPRSS2 (H296A, S441A and GS-AA) proteins were predominantly expressed as full-length protein products (∼54kDa) and lacked autocleavage products (Figure 3B, Lanes 3, 4 and 5 respectively). As expected, truncated TMPRSS2 proteins comprising LDLRA and LDLR-SRCR domains were detected as 24kDa and 37kDa products respectively (Figure 3B, Lanes 6 and 7 respectively).

To confirm the maintenance of normal cellular function upon expression of TMPRSS2 and its mutant derivatives, we infected 293T cells expressing full length TMPRSS2, its catalytically inactive mutants (H296A, S441A and GS-AA), and its domains (LDLRA, LDLRA-SRCR) with pseudoviruses bearing vesicular stomatitis virus (VSV) glycoprotein (VSV-G). VSV-G mainly binds low-density lipoprotein receptor (LDLR), but also uses other members of LDLR family as alternative receptors to gain entry (22). Following binding, VSV-G undergoes entry via clathrin-mediated endocytic pathway (23–25).VSV pseudovirus infection remained sensitive to endosomal acidification inhibitors (chloroquine, NH_4_Cl) but not TMPRSS2 protease inhibitor (camostat mesylate) (Figure 3C). Compared to VSV pseudovirus infection alone, VSV pseudovirus infection in the presence of chloroquine displayed 21-38-fold reduction whereas VSV infection in the presence of NH_4_Cl displayed 300-625-fold reduction in infectivity (Figure 3C). We observed no significant difference in VSV pseudovirus infection levels in 293T cells expressing TMPRSS2 and its mutant proteins, excluding any non-specific effects of TMPRSS2 and its mutants on cellular protein expression.

To confirm lack of TMPRSS2 catalytic activity due to H296A, S441A and GS-AA substitutions in SP domain or absence of SP domain (LDLRA, LDLRA-SRCR), we infected 293T-ACE2 cells expressing full length TMPRSS2, its catalytically inactive mutants (H296A, S441A and GS-AA) and its individual domains (LDLRA, LDLRA-SRCR) with SARS-CoV-2 pseudovirus. SARS-CoV-2 pseudovirus displayed 4-fold higher level of infection in cells expressing ACE2 and TMPRSS2 compared to cells expressing only ACE2 and remained sensitive to camostat mesylate inhibition as well as endosomal acidification inhibition, confirming SARS-CoV-2 entry by cell surface and endosomal routes (Figure 3D). SARS-CoV-2 pseudovirus infection levels in cells expressing ACE2 and various TMPRSS2 domains (LDLRA, LDLRA-SRCR) or catalytically inactive TMPRSS2 (H296A, S441A and GS-AA), was similar to that of 293T-ACE2 cells and sensitive to endosomal acidification inhibitors (chloroquine, NH_4_Cl). In the presence of chloroquine and NH_4_Cl, SARS-CoV-2 infection levels reduced 18-46-fold and 24-42-fold, respectively, in the above treatments confirming that TMPRSS2 domains (LDLRA, LDLRA-SRCR) or catalytically inactive TMPRSS2 (H296A, S441A and GS-AA) constructs lacking SP activity cannot enhance SARS-CoV-2 entry via cell surface route (Figure 3D).

We next examined whether HCoV-HKU1 pseudovirus infection is independent of TMPRSS2 SP activity. We observed HCoV-HKU1 pseudovirus infection in cells expressing full length TMPRSS2 as well as catalytically inactive TMPRSS2 (H296A, S441A and GS-AA) with pseudovirus entry being sensitive to endosomal acidification inhibitors (chloroquine, NH_4_Cl) but not TMPRSS2 SP activity inhibitor (camostat mesylate) (Figure 3E). Compared to HCoV-HKU1 pseudovirus infection without inhibitors, HCoV-HKU1 pseudovirus infection in the presence of chloroquine and NH_4_Cl displayed 6-12-fold and 5-14-fold reduction, respectively. Cells expressing TMPRSS2 domains (LDLRA, LDLRA-SRCR) failed to support HCoV-HKU1 pseudovirus infection. Based on these results, we conclude that, HKU1 spike-mediated entry requires TMPRSS2 and its SP domain but not its SP activity. TMPRSS2 facilitates HKU1 pseudovirus entry via endosomal route unlike its role in SARS-CoV-2 pseudovirus entry.

To verify cell surface expression of TMPRSS2 and its mutants, we biotinylated cell membrane proteins of 293T cells expressing TMPRSS2, catalytically inactive TMPRSS2 (H296A, S441A and GS-AA) and individual domains (LDLRA, LDLRA-SRCR) followed by immunoprecipitation by streptavidin resin and immunoblotted for N-terminal Flag tag. Consistent with previous findings, we were unable to detect full length TMPRSS2 on cell surface possibly due to ectodomain shedding caused by its autoactivation (17, 19). However, we were able to detect three N-terminal cleaved products (∼21, 25, 27kDa respectively) on cell surface (Supplementary figure 4). In line with previous study findings, we detected full length catalytically inactive TMPRSS2 mutant proteins (H296A, S441A and GS-AA) and TMPRSS2 domains (LDLRA-SRCR) sized ∼54kDa and ∼37kDa respectively (17, 19). As a positive control, cell surface expression of ACE2 was also confirmed. These findings further confirm the requirement of full length TMPRSS2 protein comprising SP domain but not its catalytic activity for HCoV-HKU1 pseudovirus infection.

### HCoV-HKU1 spike binds TMPRSS2

As HCoV-HKU1 pseudovirus infection is facilitated by full-length TMPRSS2 comprising either catalytically active or inactive SP domain, we next examined whether HCoV-HKU1 spike binds catalytically active or inactive TMPRSS2 by performing cell surface co-immune precipitation experiments. 293T target cells expressing HCoV-HKU1 spike and 293T effector cells expressing Flag-tagged TMPRSS2 proteins or individual domains described above were co-incubated prior to immunoprecipitation of TMPRSS2 proteins and individual domains with anti-Flag Ab followed by probing for HCoV-HKU1 spike binding. Effector cells expressing either full length TMPRSS2 or catalytically inactive TMPRSS2 (H296A, S441A and GS-AA) but not LDLRA or LDLRA-SRCR domains were able to pull down full length HCoV-HKU1 spike (Figure 4A). As a positive control, ACE2 but not TMPRSS2-S441A pulled down SARS-CoV-2 ΔPRRA spike (Supplementary figure 5). We used SARS-CoV-2 ΔPRRA spike to circumvent S1 subunit loss due to shedding upon ACE2 binding. Altogether, these results indicate that full length TMPRSS2 comprising either catalytically active or inactive SP domain binds HCoV-HKU1 spike to facilitate virus entry.

**Figure 4.**
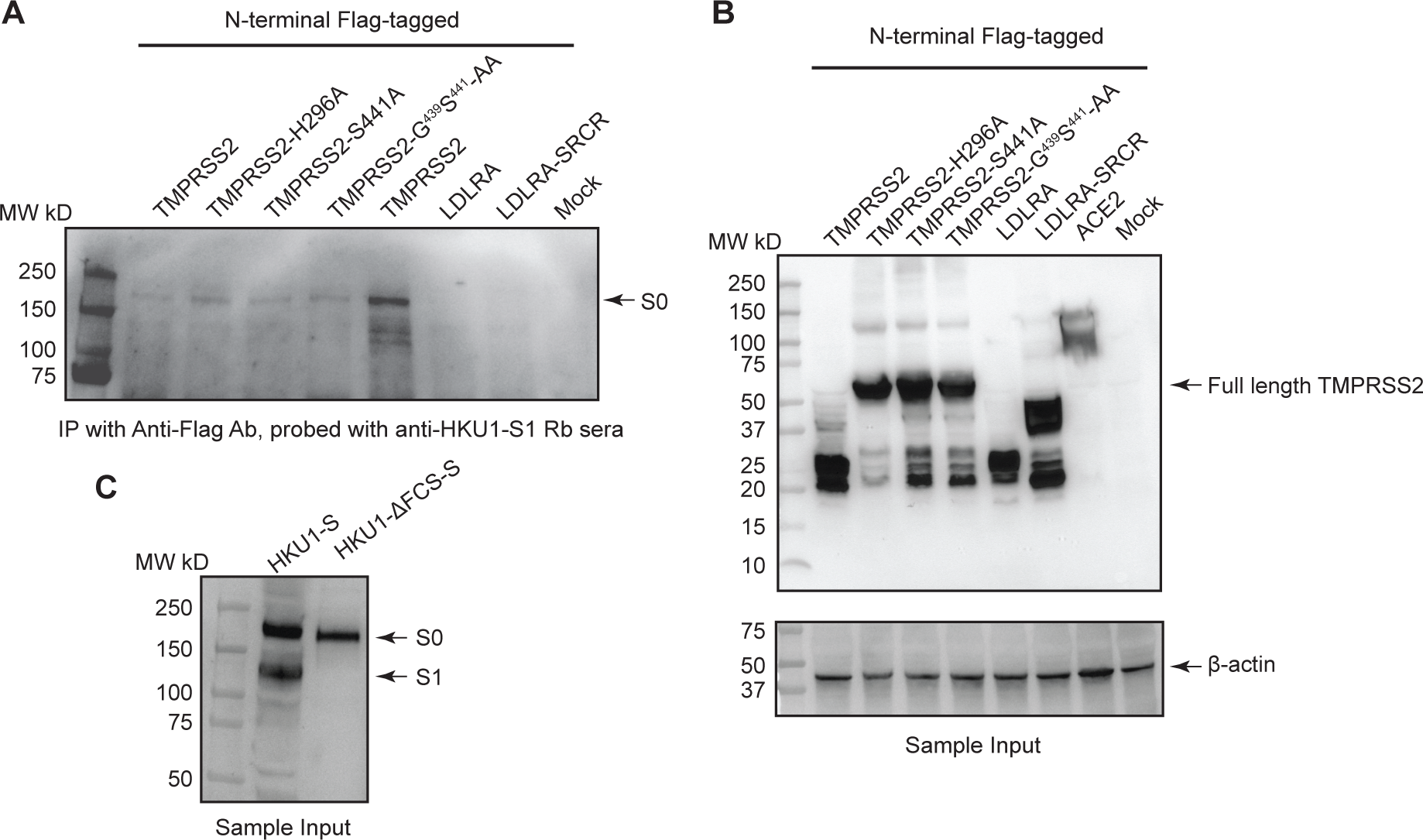
HCoV-HKU1 full length spike binds to TMPRSS2. (A) Western blotting of HCoV-HKU1 spike in Anti-Flag Ab pull downs immunoprecipitated by indicated TMPRSS2 proteins with a Flag tag at the N-terminus. HCoV-HKU1 spike is probed using polyclonal serum. (B) Sample input employed for immunoprecipitation. TMPRSS2 proteins Flag tagged in the N-terminus are probed using anti-Flag antibody. Β-actin is loading control. (C) HCoV-HKU1 spike is expressed as both full length S0 and furin processed S1 forms. HCoV-HKU1-ΔFCS, which expresses only the full length S0 form is shown for size comparison.

## Discussion

In the present study, we investigated entry mechanisms of HCoV-NL63, -229E and -HKU1 spike proteins to address some unresolved issues from prior studies as discussed below. To focus on spike-mediated entry, we generated pseudoviruses and established cells stably expressing ACE2/APN or TMPRSS2 that support high-level infection of 229E and HKU1 pseudoviruses, respectively.

In 293T-ACE2 cells, we demonstrated that HCoV-NL63 pseudoviruses followed endosomal route of entry, consistent with a previous report where authentic HCoV-NL63 was shown to be sensitive to endosomal acidification inhibitors (NH_4_Cl and bafilomycin A) in LLC-Mk2 cells (26, 27). In contrast, endosomal acidification inhibitors failed to inhibit authentic HCoV-NL63 entry in human airway epithelial (HAE) cultures, leading to a proposal that HCoV-NL63 may undergo alternate routes of entry that may involve using TMPRSS2 in HAE cells (26). Although, TMPRSS2 was shown to support or enhance entry of multiple HCoVs including SARS-CoV-2 (28–31), we found no significant enhancement of HCoV-NL63 pseudovirus infection in 293T-ACE2-TMPRSS2 cells compared to 293T-ACE2. HCoV-NL63 pseudovirus infection also remained insensitive to TMPRSS2 inhibitor camostat as shown previously (26).

Similarly, HCoV-229E pseudovirus also entered 293T-APN cells via endosomal route. This finding agrees with previous findings where authentic HCoV-229E entry was demonstrated to occur via caveolin-mediated endocytosis in human fibroblasts (32). However, HCoV-229E pseudovirus entry was shown to occur via Cathepsin L (CatL)-dependent and -independent routes in cells expressing APN and APN+TMPRSS2 respectively (33). In 293T-APN cells, coexpression of TMPRSS2 or TMPRSS11D (Human Airway Trypsin or HAT) enhanced HCoV-229E pseudovirus entry via CatL-independent route (33).

We also investigated furin-cleavage requirements for spike-mediated entry. The predicted furin cleavage sites in the S1/S2 junctions of HCoV-NL63 and -229E spike proteins do not conform to standard FCS sequence pattern (R-X-[K/R]-R); therefore, modification of predicted FCS in S1/S2 junctions did not affect spike processing (Figures 1A-D). In contrast, the FCS in S1/S2 junction of HCoV-HKU1 spike conformed to standard FCS sequence pattern (R-X-[K/R]-R) and led to efficient furin processing of spike into S1 and S2 subunit (Figures 1F). Modification of predicted FCS abolished furin processing of HCoV-HKU1 spike, which possibly prevented spontaneous shedding of the S1 subunit leading to enhanced spike-mediated infectivity (Figures 1E). Moreover, our infectivity results led us to propose that predicted FCS in HCoV-NL63 and -229E spikes may be required to maintain overall structure of spike proteins rather than furin processing function. Previous studies demonstrated considerable sequence variation and deletion between the GxCx motif and the conserved S2 nonamer in the uncleaved spike glycoproteins of group 1 HCoVs including HCoV-NL63 and -229E (34) as well as CatL requirement for HCoV-229E endocytic entry (34). Our results agree and extend previous findings that one or two basic residues in the S1/S2 junction after GxCx motif do not facilitate furin processing of HCoV-229E and -NL63 spikes respectively.

For HCoV-HKU1, we refined the role of TMPRSS2 in supporting virus entry. We found that HCoV-HKU1pseudovirus infection was dependent on TMPRSS2 and followed endosomal route of entry but did not require its SP catalytic activity. TMPRSS2 is N-glycosylated type II transmembrane serine protease belonging to TTSP family which comprises several other serine proteases such as hepsin, matriptase, TMPRSS11D, and corin (35–38). In human airways, TMPRSS2 has divergent roles in viral infections including processing and activation of viral glycoproteins (e.g., influenza hemagglutinin (39–41), human metapneumovirus (42), and SARS-CoV-1 spike (43)), enhancing viral entry by S2’ priming (e.g., SARS-CoV-2 (31)) and cleavage and downregulation of cellular receptors of viruses (e.g., ACE2 (44)). Typically, TMPRSS2 SP functions are limited to cell surface and early endosomes (pH∼6.8) where it promotes S2’ site cleavage and membrane fusion during SARS-CoV-2 entry (45). At pH<6.8, TMPRSS2 SP activity is expected to drop and therefore, SARS-CoV-2 entry in late endosomal/lysosomal compartments is facilitated by CatL priming of S2’ (45). In the present study, HCoV-HKU1 spike-mediated entry required acidic pH as shown by sensitivity to endosomal acidification inhibitors (chloroquine and NH_4_Cl) pointing to entry in late endosomal/lysosomal compartments where TMPRSS2 SP activity might be low. In support of this, HCoV-HKU1 spike-mediated entry is not affected upon abolishing TMPRSS2 SP activity through H296A, S441A and GS-AA substitutions and/or in the presence of TMPRSS2 protease inhibitor camostat. Through infectivity and immunoprecipitation experiments, we showed that HKU1 spike binding and entry requires SP domain but not SP activity. This finding is in line with overall receptor utilization requirements of coronavirus spike. Enzymatic inhibition of porcine APN, human dipeptidyl peptidase 4 and human ACE2 receptors has no effect on the infection of transmissible gastroenteritis virus (46), MERS-CoV (47, 48), and SARS-CoV-2 (49, 50) respectively, suggesting that while coronavirus spikes may utilize distinct receptor domains for binding, they do not require enzymatic activity of the receptor (46, 51).

While our study was under preparation, a recent study has demonstrated TMPRSS2 to be the functional receptor of HCoV-HKU1 regardless of TMPRSS2 serine protease activity, although TMPRSS2 activity was found to be required for facilitating cell-cell fusion (9). This study demonstrated HCoV-HKU1 entry in 293T cells expressing active and inactive TMPRSS2 containing R255Q and S441A single amino acid substitutions. Our results extend these findings to show that H296A, S441A and GS-AA substitutions of inactive TMPRSS2 also facilitate HCoV-HKU1 entry. Furthermore, in that study, two cathepsin inhibitors (SB412515 and E64d) and endosomal acidification inhibitor hydroxychloroquine were demonstrated to inhibit entry in inactive TMPRSS2-expressing 293T cells but not active TMPRSS2-expressing cells. This led the authors to propose that pseudoviruses expressing HKU1A spike can either fuse at the plasma membrane if the protease is active or be internalized and processed in the endosomal compartment when the protease is inactive. We show that HCoV-HKU1A spike (spike used in the current study belongs to HKU1A genotype) require endosomal acidification for efficient entry since TMPRSS2 inhibitor failed to inhibit HCoV-HKU1 pseudovirus entry in cells expressing active or inactive TMPRSS2 but endosomal acidification inhibitors did inhibit entry. However redundant roles of active and inactive TMPRSS2 in facilitating entry and cell-cell fusion cannot be excluded. While previous study demonstrated binding of soluble spike or RBD to active and inactive TMPRSS2, we showed binding of full-length spike to TMPRSS2 with an active or inactive serine protease domain.

In summary, HCoV-NL63, -229E and HKU1 pseudoviruses entered via endosomal route of entry. Whereas the predicted S1/S2 junction residues of HCoV-NL63 and -229E spike proteins are required for optimal cellular entry, multibasic residues in the S1/S2 junction of HCoV-HKU1 spike are not required for entry. Finally, HCoV-HKU1 entry via endosomal route requires cell surface TMPRSS2 and its serine protease domain, but not its serine protease activity. Limitations of our study include examining entry mechanisms in 293T cells rather than physiologically relevant target cell types and the use of lentiviral pseudoviruses instead of authentic viruses.

## Materials and Methods

### Plasmids, cell lines and inhibitors

Codon-optimized full length open reading frames of spike genes of HCoV-NL63 (Genbank Accession: YP_003767.1), HCoV-229E (Genbank Accession: NP_073551), HCoV-HKU1 (Genbank Accession: YP_173238.1, genotype 1A), and SARS-CoV-2 D614G (GISAID Accession: EPI_ISL_5851484) were synthetized by GenScript (Piscataway, NJ) and cloned into pVRC8400 or pcDNA3.1(+) vectors using standard molecular biology protocols and confirmed by Sanger sequencing. Deletions in the predicted furin cleavage sites of HCoV-NL63, -229E and -HKU1 spikes were performed using QuikChange one-step site-directed deletion mutagenesis protocol as described previously (52) and confirmed by sequencing. Human ACE2 (GenBank accession: NM_021804.2) with a C-terminal Flag tag was synthesized by GenScript (Piscataway, NJ) and pCSDest-TMPRSS2 (Addgene Plasmid # 53887) plasmid was obtained from Addgene as described previously (53). An N-terminal Flag tag was introduced after start codon in pCSDest-TMPRSS2 plasmid to create TMPRSS2-Flag-N using one-step site-directed insertion mutagenesis protocol using primers described in Table 1 (52). To abolish TMPRSS2 serine protease activity, catalytic residues (H296A, S441A) in TMPRSS2 serine protease domain were mutated to alanine in TMPRSS2-Flag-N by site-directed mutagenesis. The backbone amine groups of G439 and S441 contribute to the oxyanion hole formation which help to activate and stabilize the carbonyl of the scissile bond during TMPRSS2-mediated proteolysis (21). Double alanine mutations G^439^S^441^-AA in the G^439^DS^441^GG motif were introduced to prevent oxyanion hole formation and further prevent TMPRSS2 proteolytic activity (21). The Low-Density Lipoprotein Receptor A (LDLRA) and Scavenger Receptor-Cysteine Rich (SRCR) domains of TMPRSS2 (GenBank Accession: NP_001128571.1) span between 149-186 aa and 195-226 aa respectively (54). To generate plasmids expressing LDLRA and both LDLRA and SRCR domains of TMPRSS2, Y198 and R^292^I^293^ aa positions in TMPRSS2-Flag-N are mutated to one and two consecutive stop codons respectively. All single- and double-point mutations were introduced using Agilent QuikChange site-directed mutagenesis kit using primers described in Table 1.

**Table 1.**
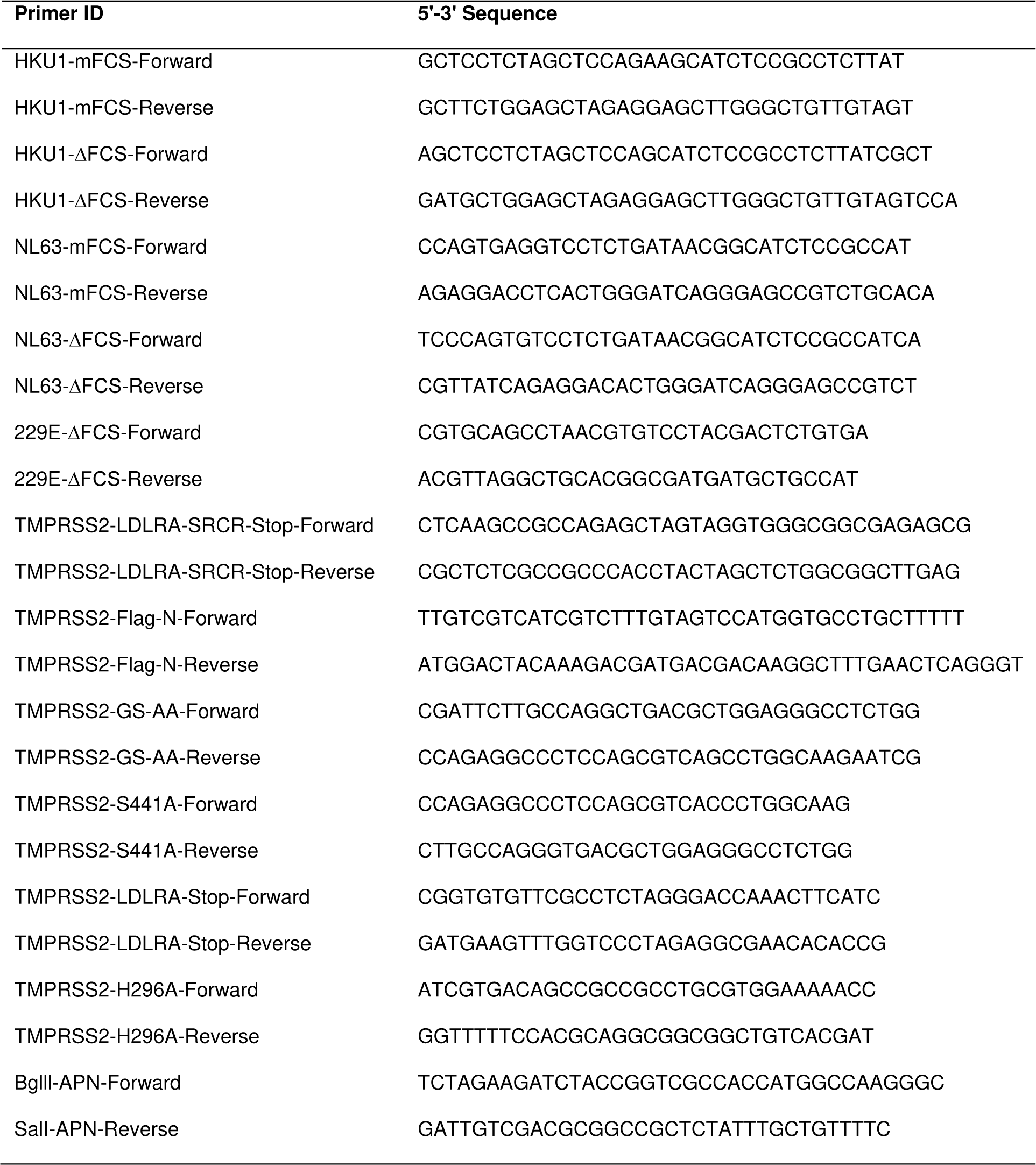
List of primers described in the study.

The HIV gag/pol (pCMVΔR8.2), pCMV/R, and Luciferase reporter (pHR’CMV-Luc) plasmids were obtained from the Vaccine Research Center (National Institutes of Health (NIH), Bethesda, MD) as described previously (55, 56). pCMV-VSV-G was obtained from Dr. Kathy Bouir, (University of California, San Diego) (12). pLenti-CMV-GFP-Puro (plasmid #17448) was obtained from Addgene. To generate pLenti-CMV-hAPN-Puro, full length open reading frame of human aminopeptidase N (hAPN) was PCR amplified from pVRC-hAPN using primers flanking BgllI and SalI sites, cloned into pCR™TOPO2.1 using TOPO™ TA Cloning™ Kit (Invitrogen) and subcloned into pLenti-CMV-GFP-Puro (Addgene Plasmid #17448). pHAGE2-EF1aInt-TMPRSS2-IRES-mCherry plasmid was kindly provided by Dr. Jesse Bloom (Fred Hutchinson Cancer Center, Seattle, WA) (57). HEK293T-ACE2 cells stably expressing ACE2 were obtained through BEI Resources (BEI Resources, Cat no: NR-52511). HEK293T-ACE2-TMPRSS2 cells stably expressing both ACE2 and TMPRSS2 were generated as described previously (BEI Resources, Cat no: NR-55293) (12). Cell lines including 293T, A549, Calu-3, HCT-8, Caco-2, Huh-7, BHK21, U87, rhabdomyosarcoma (RD) and Vero cells were maintained at 37°C in Dulbecco’s modified eagle medium (DMEM) supplemented with high glucose, L-Glutamine, minimal essential media (MEM) non-essential amino acids, penicillin/streptomycin and 10% fetal bovine serum (FBS). Chemical inhibitors, camostat mesylate (TMPRSS2 inhibitor; Cat no: SML0057), chloroquine (endosomal acidification inhibitor; Cat no: 50-63-5) and ammonium chloride (endosomal acidification inhibitor; Cat no: 12125-02-9) were obtained from Millipore Sigma.

### Antibodies and sera

Rabbit sera against HCoV-NL63 spike was generated by priming New Zealand white rabbit intramuscularly with 45 µg purified His-tagged NL63 Spike-S1-RBD (Genscript) complexed with Freund’s complete adjuvant followed by 3 boosts with 45 µg purified His-tagged HCoV-NL63 Spike-S1-RBD (first boost) and subsequent boosts with 25 µg purified HCoV-NL63 stabilized spike (provided by Christopher Broder) mixed in Freund’s incomplete adjuvant, at 3-4-week intervals. Control rabbit sera was obtained from same rabbit prior to immunization. Rabbit sera against HCoV-229E spike was generated by priming New Zealand white rabbit intramuscularly with 59 µg purified recombinant His-tagged HCoV-229E Spike-S1 subunit (Sino Biological Cat No: 40601-V08H) complexed with Freund’s complete adjuvant followed by three boosts with 59 µg purified His-tagged HCoV-229E Spike-S1 subunit (first boost) and subsequent boosts with 25 µg purified HCoV-229E stabilized spike (provided by Christopher Broder) mixed in Freund’s incomplete adjuvant, at 3-4-week intervals. Control rabbit sera was obtained from same rabbit prior to immunization. Rabbit sera against HCoV-HKU1 spike was generated by priming New Zealand white rabbit intramuscularly with 59 µg purified recombinant His-tagged HCoV-HKU1 Spike-S1 subunit (Sino Biological Cat No: 40021-V08H) complexed with Freund’s complete adjuvant followed by three boosts with 59 µg purified His-tagged HCoV-HKU1 Spike-S1 subunit mixed with Freund’s incomplete adjuvant, at 3-4-week intervals. Control rabbit sera was obtained from same rabbit prior to immunization. All serum samples were heated to 56°C for 30 min to inactivate complement.

Mouse monoclonal Ab (ID: 9D3F3) against HCoV-NL63 S1 subunit and mouse monoclonal Ab (ID: 24C9F7) against HCoV-229E S1 subunit were generated by Genscript. Rabbit polyclonal Ab against SARS-CoV-2 RBD was obtained from Sino Biological (Cat No: 40592-T62). Mouse monoclonal Ab against the Flag tag, THE™ DYKDDDDK Tag Antibody, was obtained from Gencript (Cat no: A00187). Mouse monoclonal Ab against β-actin, THE™ beta Actin Antibody [HRP], was obtained from Gencript (Cat no: A00730). HIV-1 p24 hybridoma (183-H12-5C) was obtained through the AIDS Research and Reference Reagent Program, Division of AIDS, NIAID, NIH and kindly provided by Dr. Bruce Chesebro.

### Pseudovirus production and neutralization

Pseudoviruses bearing the desired spike glycoprotein of HCoV-NL63, -229E, -HKU1 and SARS-CoV-2 and carrying a firefly luciferase (FLuc) reporter gene were produced by cotransfecting HEK-293T (ATCC, Cat no: CRL-11268) cells with 5 μg of pCMVΔR8.2, 5 μg of pHR’CMVLuc and 0.5 μg of pVRC spike or its corresponding mutant expression plasmids. Pseudovirus supernatants were collected approximately 48 h post-transfection, filtered through a 0.45 μm low protein binding filter, and used immediately or stored at -80°C. Pseudovirus titers were measured by infecting target cells for 48 h prior to measuring luciferase activity (luciferase assay reagent, Promega, Madison, WI), as described previously (58). Pseudovirus titers were expressed as relative luminescence unit per milliliter of pseudovirus supernatants (RLU/ml). Pseudovirus neutralization assays were performed in 96-well plates as previously described (12). Pseudoviruses with titers of approximately 10^6^ RLU/mL of luciferase activity were incubated with serially diluted rabbit sera for two hours at 37°C prior to inoculation onto the plates that were pre-seeded one day earlier with 3.0 × 10^5^ cells/ml. Pseudovirus infectivity was determined 48 h post inoculation for luciferase activity by luciferase assay reagent. The inverse of the sera dilutions causing a 50% reduction of RLU compared to control was reported as the neutralization titer (ID50). Titers were calculated using a nonlinear regression curve fit (GraphPad Prism Software Inc., La Jolla, CA, USA). The mean titer from at least two independent experiments each with intra-assay duplicates was reported as the final titer. For experiments involving camostat mesylate, chloroquine and NH_4_Cl inhibitors, each target cell type was pretreated with inhibitor for 2 h before pseudovirus infection in the presence of the respective inhibitor as described previously (12).

### Establishment of stable cell lines and transiently transfected cells

For establishing 293T-ACE2-APN, and 293T-TMPRSS2 cells that stably express APN and TMPRSS2 proteins respectively, VSV-G-pseudotyped lentiviruses carrying the human APN or TMPRSS2 gene were generated by co-transfecting 293T cells with 5 μg of pLenti-CMV-hAPN-Puro or pHAGE2-EF1aInt-TMPRSS2-IRES-mCherry, 5 μg of pCMVΔ8.2, and 0.5 μg of pCMV-VSV-G plasmids. Packaged lentivirus was harvested 48h post transfection and used to transduce 293T-ACE2 (BEI Resources: NR-52511) and 293T cells in the presence of 10 μg/mL polybrene by spinoculation at 2000 x g for 30 min at room temperature (59). The medium was replaced 24 hours post transduction and expanded for another 48 hours. The selection marker for 293T-TMPRSS2 cells is mCherry and therefore, bulk transduced population was single-cell sorted based on high mCherry positivity into clear bottomed 96 well plates by flow cytometry on a BD FACSAria II Cell Sorter as described previously (12). We expanded one clone (H10) that displayed high-levels of infection with HCoV-HKU1 pseudovirus and stably expressed TMPRSS2 at least until 20 passages. The selection marker for 293T-ACE2-APN cells is puromycin, and therefore, bulk transduced population was selected with puromycin (2 µG/mL) followed by the expansion of single cell clones obtained through limiting dilution cloning method. Expanded clone of 293T-ACE2-APN-Puro cells supported high levels of infection with HCoV-229E pseudovirus and remained stable at least until 50 passages when maintained under 1-2 µG/mL puromycin selection.

We assessed HCoV-NL63, -229E and -HKU1 pseudoviruses for infectivity in stable and continuous cell lines for their ability to support pseudovirus infection. Previous studies reported that 293T, 293T-ACE2, 293T-ACE2-TMPRSS2, Huh-7, and Huh-7-ACE2-TMPRSS2 cells supported infection of HCoV-NL63 lentiviral-based pseudoviruses and coexpression of TMPRSS2 along with ACE2 led to several fold enhancement in infection titers (10, 60). For HCoV-NL63 pseudovirus, we found that cell lines expressing ACE2 receptor, 293T-ACE2, 293T-ACE2-TMPRSS2 and 293T-ACE2-APN supported 40-fold, 33-fold and 11-fold higher levels of infection compared to background levels of infection in 293T cells (10^4^ RLU/ml) (Supplementary Figure 1A). None of other cell lines supported infection by HCoV-NL63 pseudoviruses. Cell types lacking ACE2 receptor (293T, A549) did not support HCoV-NL63 pseudovirus infection and coexpression of TMPRSS2 along with ACE2 in 293T-ACE2-TMPRSS2 cells did not necessarily enhance infection compared to 293T-ACE2 cells expressing ACE2 receptor itself (Supplementary Figure 1A).

For HCoV-229E pseudoviruses, except 293T-ACE2-APN cells expressing hAPN receptor, none of other cell lines supported infection (Supplementary Figure 1B). Stable expression of hAPN resulted in 3800-fold higher levels of infection compared to 293T cells with no expression of hAPN.

For HCoV-HKU1 pseudoviruses, cell lines with stable expression of TMPRSS2, 293T-TMPRSS2 and 293T-ACE-TMPRSS2 supported 207-fold and 27-fold higher levels of infection compared to 293T cells with background infection levels (Supplementary Figure 1C). None of other cell lines supported HCoV-HKU1 pseudovirus infection presumably due to lack of TMPRSS2 expression. These findings are in line with previous findings where TMPRSS2 expression in 293T cells supported HCoV-HKU1 pseudovirus infection (10). As a control, SARS-CoV-2 pseudovirus infection was supported by cell lines expressing ACE2 including 293T-ACE2, 293T-ACE2-TMPRSS2 and 293T-ACE2-APN and coexpression of TMPRSS2 led to 10-fold higher infection (Supplementary Figure 1D). Since 293T-ACE2, 293T-ACE2-APN and 293T-TMPRSS2 cells supported optimal levels of infection by HCoV-NL63, -229E and - HKU1 pseudoviruses respectively, all the experiments reported in the study were performed in these cells.

For transient transfections, 4 μg TMPRSS2-Flag-N or its point mutations’ encoding plasmids described above were transfected alone or in combination with 4μg ACE2-Flag into 293T cells using FuGene transfection reagent (Promega). Cells were infected with HCoV-HKU1, VSV-G or SARS-CoV-2 pseudoviruses and quantified for luciferase activity 48 hours post infection. Protein expression was confirmed by Western Blotting.

### Cell Surface Co-immune precipitation

293T target cells were transfected with a plasmid encoding HCoV-HKU1 or SARS-CoV-2 ΔPRRA spike. 293T effector cells were separately transfected with plasmids encoding Flag-tagged ACE2, TMPRSS2 and TMPRSS2 mutants. Two days post transfection, target and effector cells were dissociated by cell dissociation solution (Thermo Scientific™), washed with cell culture medium and co-incubated in equal numbers for two hours by nutation. Coincubated cells were spun at 800 *x g*, gently washed with cell culture medium three times, and lysed in NP-40 lysis buffer containing Halt™ Protease and Phosphatase Inhibitor Cocktail (Thermo Scientific™). Cell lysates were immunoprecipitated for an hour using Pierce™ Protein G Magnetic Beads that are preconjugated with THE™ DYKDDDDK Tag antibody directed against Flag tag. Immune complexes were washed three times to exclude any non-specific binding followed by elution in Laemmli buffer (Biorad) containing 1 mM DTT and Western Blotting of eluates for HCoV-HKU1 or SARS-CoV-2 ΔPRRA spike.

### Cell surface protein biotinylation

HEK293T cells expressing TMPRSS2 proteins or its domains in 6-well plates were washed with PBS and incubated with EZ-Link™ Sulfo-N-Hydroxy Succinimide– SS-Biotin (0.25 mg/ml, 1 ml/well) (Thermo Fisher Scientific; Cat No: A44390) label for 20 min at room temperature. The labeling solution is then removed, and cells are washed thrice with 2 ml ice-cold BupH™ Tris-buffered saline solution to stop the reaction. Cells were then lysed, and Biotin-labeled proteins were precipitated with NeutrAvidin agarose beads (Thermo Fisher) for an hour followed by three wash steps in wash buffer, elution, and Western Blotting of eluted proteins.

### Western Blotting

HCoV-NL63, -229E, -HKU1 and SARS-CoV-2 pseudoviruses were resolved by 4-20% SDS-PAGE and probed for spike protein using mouse monoclonal Ab 9D3F3, mouse monoclonal Ab 24C9F7, rabbit polyclonal sera, and rabbit polyclonal Ab respectively. Coimmuneprecipitation eluates were resolved by 4-20% SDS-PAGE and probed for spike using rabbit sera, and rabbit polyclonal Ab for HCoV-HKU1 and SARS-CoV-2 ΔPRRA spike respectively. Biotin-labeled TMPRSS2 proteins and its domains were probed using Flag Ab. Transfected cell lysates were prepared in NP-40 lysis buffer containing Halt™ Protease Inhibitor Cocktail (Thermo Scientific™), resolved by 4-20% SDS-PAGE and probed using suitable primary Abs. Membranes probed with primary Abs were detected using HRP-conjugated species-specific secondary antibodies and luminol-based chemiluminescence (Sera care, Milford, MA).

### Statistical analyses

Two-group comparisons (Mann-Whitney test) were analyzed using GraphPad Prism software. *P values ≤ 0.05* were considered statistically significant.

## ACKNOWLEDGMENTS

We would like to thank Shuang Tang (FDA) and Hailun Ma (FDA) for the critical review of the manuscript. We do not have commercial or other associations or competing interests that might pose a conflict of interest.

Financial support was provided by the U.S. Food and Drug Administration.

## References

1. Shah MM, Winn A, Dahl RM, Kniss KL, Silk BJ, Killerby ME. 2022. Seasonality of Common Human Coronaviruses, United States, 2014-2021(1). Emerg Infect Dis 28:1970–1976.

2. Edridge AWD, Kaczorowska J, Hoste ACR, Bakker M, Klein M, Loens K, Jebbink MF, Matser A, Kinsella CM, Rueda P, Ieven M, Goossens H, Prins M, Sastre P, Deijs M, van der Hoek L. 2020. Seasonal coronavirus protective immunity is short-lasting. Nature Medicine 26:1691–1693.

3. Callow KA, Parry HF, Sergeant M, Tyrrell DA. 1990. The time course of the immune response to experimental coronavirus infection of man. Epidemiol Infect 105:435–46.

4. Kistler KE, Bedford T. 2021. Evidence for adaptive evolution in the receptor-binding domain of seasonal coronaviruses OC43 and 229e. eLife 10:e64509.

5. Eguia RT, Crawford KHD, Stevens-Ayers T, Kelnhofer-Millevolte L, Greninger AL, Englund JA, Boeckh MJ, Bloom JD. 2021. A human coronavirus evolves antigenically to escape antibody immunity. PLOS Pathogens 17:e1009453.

6. Wang C, Hesketh EL, Shamorkina TM, Li W, Franken PJ, Drabek D, van Haperen R, Townend S, van Kuppeveld FJM, Grosveld F, Ranson NA, Snijder J, de Groot RJ, Hurdiss DL, Bosch B-J. 2022. Antigenic structure of the human coronavirus OC43 spike reveals exposed and occluded neutralizing epitopes. Nature Communications 13:2921.

7. Cantoni D, Siracusano G, Mayora-Neto M, Pastori C, Fantoni T, Lytras S, Di Genova C, Hughes J, On Behalf Of The Ambulatorio Medico San Luca Villanuova G, Lopalco L, Temperton N. 2022. Analysis of Antibody Neutralisation Activity against SARS-CoV-2 Variants and Seasonal Human Coronaviruses NL63, HKU1, and 229E Induced by Three Different COVID-19 Vaccine Platforms. Vaccines (Basel) 11.

8. Chen Y, Zhao X, Zhou H, Zhu H, Jiang S, Wang P. 2023. Broadly neutralizing antibodies to SARS-CoV-2 and other human coronaviruses. Nature Reviews Immunology 23:189–199.

9. Saunders N, Fernandez I, Planchais C, Michel V, Rajah MM, Baquero Salazar E, Postal J, Porrot F, Guivel-Benhassine F, Blanc C, Chauveau-Le Friec G, Martin A, Grzelak L, Oktavia RM, Meola A, Ahouzi O, Hoover-Watson H, Prot M, Delaune D, Cornelissen M, Deijs M, Meriaux V, Mouquet H, Simon-Lorière E, van der Hoek L, Lafaye P, Rey F, Buchrieser J, Schwartz O. 2023. TMPRSS2 is a functional receptor for human coronavirus HKU1. Nature 624:207–214.

10. Sampson AT, Heeney J, Cantoni D, Ferrari M, Sans MS, George C, Di Genova C, Mayora Neto M, Einhauser S, Asbach B, Wagner R, Baxendale H, Temperton N, Carnell G. 2021. Coronavirus Pseudotypes for All Circulating Human Coronaviruses for Quantification of Cross-Neutralizing Antibody Responses. Viruses 13.

11. Crawford KHD, Eguia R, Dingens AS, Loes AN, Malone KD, Wolf CR, Chu HY, Tortorici MA, Veesler D, Murphy M, Pettie D, King NP, Balazs AB, Bloom JD. 2020. Protocol and Reagents for Pseudotyping Lentiviral Particles with SARS-CoV-2 Spike Protein for Neutralization Assays. Viruses 12:513.

12. Neerukonda SN, Vassell R, Herrup R, Liu S, Wang T, Takeda K, Yang Y, Lin T-L, Wang W, Weiss CD. 2021. Establishment of a well-characterized SARS-CoV-2 lentiviral pseudovirus neutralization assay using 293T cells with stable expression of ACE2 and TMPRSS2. PLOS ONE 16:e0248348.

13. Coutard B, Valle C, de Lamballerie X, Canard B, Seidah NG, Decroly E. 2020. The spike glycoprotein of the new coronavirus 2019-nCoV contains a furin-like cleavage site absent in CoV of the same clade. Antiviral Res 176:104742.

14. Bestle D, Heindl MR, Limburg H, Van Lam van T, Pilgram O, Moulton H, Stein DA, Hardes K, Eickmann M, Dolnik O, Rohde C, Klenk HD, Garten W, Steinmetzer T, Böttcher-Friebertshäuser E. 2020. TMPRSS2 and furin are both essential for proteolytic activation of SARS-CoV-2 in human airway cells. Life Sci Alliance 3.

15. Tian S. 2009. A 20 Residues Motif Delineates the Furin Cleavage Site and its Physical Properties May Influence Viral Fusion. Biochemistry Insights 2:BCI.S2049.

16. Tian S, Huang Q, Fang Y, Wu J. 2011. FurinDB: A Database of 20-Residue Furin Cleavage Site Motifs, Substrates and Their Associated Drugs. International Journal of Molecular Sciences 12:1060–1065.

17. Afar DE, Vivanco I, Hubert RS, Kuo J, Chen E, Saffran DC, Raitano AB, Jakobovits A. 2001. Catalytic cleavage of the androgen-regulated TMPRSS2 protease results in its secretion by prostate and prostate cancer epithelia. Cancer Res 61:1686–92.

18. Fraser BJ, Beldar S, Seitova A, Hutchinson A, Mannar D, Li Y, Kwon D, Tan R, Wilson RP, Leopold K, Subramaniam S, Halabelian L, Arrowsmith CH, Bénard F. 2022. Structure and activity of human TMPRSS2 protease implicated in SARS-CoV-2 activation. Nature Chemical Biology 18:963–971.

19. Zhang Y, Sun S, Du C, Hu K, Zhang C, Liu M, Wu Q, Dong N. 2022. Transmembrane serine protease TMPRSS2 implicated in SARS-CoV-2 infection is autoactivated intracellularly and requires N-glycosylation for regulation. Journal of Biological Chemistry 298.

20. Böttcher E, Matrosovich T, Beyerle M, Klenk HD, Garten W, Matrosovich M. 2006. Proteolytic activation of influenza viruses by serine proteases TMPRSS2 and HAT from human airway epithelium. J Virol 80:9896–8.

21. Hempel T, Raich L, Olsson S, Azouz NP, Klingler AM, Hoffmann M, Pöhlmann S, Rothenberg ME, Noé F. 2021. Molecular mechanism of inhibiting the SARS-CoV-2 cell entry facilitator TMPRSS2 with camostat and nafamostat. Chemical Science 12:983–992.

22. Finkelshtein D, Werman A, Novick D, Barak S, Rubinstein M. 2013. LDL receptor and its family members serve as the cellular receptors for vesicular stomatitis virus. Proceedings of the National Academy of Sciences 110:7306–7311.

23. Cureton DK, Massol RH, Saffarian S, Kirchhausen TL, Whelan SP. 2009. Vesicular stomatitis virus enters cells through vesicles incompletely coated with clathrin that depend upon actin for internalization. PLoS Pathog 5:e1000394.

24. Johannsdottir HK, Mancini R, Kartenbeck J, Amato L, Helenius A. 2009. Host cell factors and functions involved in vesicular stomatitis virus entry. J Virol 83:440–53.

25. Roche S, Bressanelli S, Rey FA, Gaudin Y. 2006. Crystal structure of the low-pH form of the vesicular stomatitis virus glycoprotein G. Science 313:187–91.

26. Milewska A, Nowak P, Owczarek K, Szczepanski A, Zarebski M, Hoang A, Berniak K, Wojarski J, Zeglen S, Baster Z, Rajfur Z, Pyrc K. 2018. Entry of Human Coronavirus NL63 into the Cell. J Virol 92.

27. Huang IC, Bosch BJ, Li F, Li W, Lee KH, Ghiran S, Vasilieva N, Dermody TS, Harrison SC, Dormitzer PR, Farzan M, Rottier PJM, Choe H. 2006. SARS Coronavirus, but Not Human Coronavirus NL63, Utilizes Cathepsin L to Infect ACE2-expressing Cells*. Journal of Biological Chemistry 281:3198–3203.

28. Park JE, Li K, Barlan A, Fehr AR, Perlman S, McCray PB, Jr., Gallagher T. 2016. Proteolytic processing of Middle East respiratory syndrome coronavirus spikes expands virus tropism. Proc Natl Acad Sci U S A 113:12262–12267.

29. Reinke LM, Spiegel M, Plegge T, Hartleib A, Nehlmeier I, Gierer S, Hoffmann M, Hofmann-Winkler H, Winkler M, Pöhlmann S. 2017. Different residues in the SARS-CoV spike protein determine cleavage and activation by the host cell protease TMPRSS2. PLoS One 12:e0179177.

30. Shirato K, Kanou K, Kawase M, Matsuyama S. 2017. Clinical Isolates of Human Coronavirus 229E Bypass the Endosome for Cell Entry. J Virol 91.

31. Hoffmann M, Kleine-Weber H, Schroeder S, Krüger N, Herrler T, Erichsen S, Schiergens TS, Herrler G, Wu N-H, Nitsche A, Müller MA, Drosten C, Pöhlmann S. 2020. SARS-CoV-2 Cell Entry Depends on ACE2 and TMPRSS2 and Is Blocked by a Clinically Proven Protease Inhibitor. Cell 181:271–280.e8.

32. Nomura R, Kiyota A, Suzaki E, Kataoka K, Ohe Y, Miyamoto K, Senda T, Fujimoto T. 2004. Human coronavirus 229E binds to CD13 in rafts and enters the cell through caveolae. J Virol 78:8701–8.

33. Bertram S, Dijkman R, Habjan M, Heurich A, Gierer S, Glowacka I, Welsch K, Winkler M, Schneider H, Hofmann-Winkler H, Thiel V, Pöhlmann S. 2013. TMPRSS2 activates the human coronavirus 229E for cathepsin-independent host cell entry and is expressed in viral target cells in the respiratory epithelium. J Virol 87:6150–60.

34. Kawase M, Shirato K, Matsuyama S, Taguchi F. 2009. Protease-mediated entry via the endosome of human coronavirus 229E. J Virol 83:712–21.

35. Li S, Peng J, Wang H, Zhang W, Brown JM, Zhou Y, Wu Q. 2020. Hepsin enhances liver metabolism and inhibits adipocyte browning in mice. Proceedings of the National Academy of Sciences 117:12359–12367.

36. Antalis TM, Bugge TH, Wu Q. 2011. Chapter 1 - Membrane-Anchored Serine Proteases in Health and Disease, p 1-50. *In* Di Cera E (ed), Progress in Molecular Biology and Translational Science, vol 99. Academic Press.

37. Bugge TH, Antalis TM, Wu Q. 2009. Type II Transmembrane Serine Proteases*. Journal of Biological Chemistry 284:23177–23181.

38. Paoloni-Giacobino A, Chen H, Peitsch MC, Rossier C, Antonarakis SE. 1997. Cloning of the TMPRSS2 Gene, Which Encodes a Novel Serine Protease with Transmembrane, LDLRA, and SRCR Domains and Maps to 21q22.3. Genomics 44:309–320.

39. Bertram S, Glowacka I, Blazejewska P, Soilleux E, Allen P, Danisch S, Steffen I, Choi SY, Park Y, Schneider H, Schughart K, Pohlmann S. 2010. TMPRSS2 and TMPRSS4 facilitate trypsin-independent spread of influenza virus in Caco-2 cells. J Virol 84:10016–25.

40. Bottcher E, Matrosovich T, Beyerle M, Klenk HD, Garten W, Matrosovich M. 2006. Proteolytic activation of influenza viruses by serine proteases TMPRSS2 and HAT from human airway epithelium. J Virol 80:9896–8.

41. Bottcher-Friebertshauser E, Stein DA, Klenk HD, Garten W. 2011. Inhibition of influenza virus infection in human airway cell cultures by an antisense peptide-conjugated morpholino oligomer targeting the hemagglutinin-activating protease TMPRSS2. J Virol 85:1554–62.

42. Shirogane Y, Takeda M, Iwasaki M, Ishiguro N, Takeuchi H, Nakatsu Y, Tahara M, Kikuta H, Yanagi Y. 2008. Efficient multiplication of human metapneumovirus in Vero cells expressing the transmembrane serine protease TMPRSS2. J Virol 82:8942–6.

43. Shulla A, Heald-Sargent T, Subramanya G, Zhao J, Perlman S, Gallagher T. 2011. A transmembrane serine protease is linked to the severe acute respiratory syndrome coronavirus receptor and activates virus entry. J Virol 85:873–82.

44. Glowacka I, Bertram S, Herzog P, Pfefferle S, Steffen I, Muench MO, Simmons G, Hofmann H, Kuri T, Weber F, Eichler J, Drosten C, Pöhlmann S. 2010. Differential Downregulation of ACE2 by the Spike Proteins of Severe Acute Respiratory Syndrome Coronavirus and Human Coronavirus NL63. Journal of Virology 84:1198–1205.

45. Kreutzberger AJB, Sanyal A, Saminathan A, Bloyet L-M, Stumpf S, Liu Z, Ojha R, Patjas MT, Geneid A, Scanavachi G, Doyle CA, Somerville E, Correia RBDC, Di Caprio G, Toppila-Salmi S, Mäkitie A, Kiessling V, Vapalahti O, Whelan SPJ, Balistreri G, Kirchhausen T. 2022. SARS-CoV-2 requires acidic pH to infect cells. Proceedings of the National Academy of Sciences 119:e2209514119.

46. Delmas B, Gelfi J, Kut E, Sjöström H, Noren O, Laude H. 1994. Determinants essential for the transmissible gastroenteritis virus-receptor interaction reside within a domain of aminopeptidase-N that is distinct from the enzymatic site. Journal of Virology 68:5216–5224.

47. Wang N, Shi X, Jiang L, Zhang S, Wang D, Tong P, Guo D, Fu L, Cui Y, Liu X, Arledge KC, Chen Y-H, Zhang L, Wang X. 2013. Structure of MERS-CoV spike receptor-binding domain complexed with human receptor DPP4. Cell Research 23:986–993.

48. Raj VS, Mou H, Smits SL, Dekkers DHW, Müller MA, Dijkman R, Muth D, Demmers JAA, Zaki A, Fouchier RAM, Thiel V, Drosten C, Rottier PJM, Osterhaus ADME, Bosch BJ, Haagmans BL. 2013. Dipeptidyl peptidase 4 is a functional receptor for the emerging human coronavirus-EMC. Nature 495:251–254.

49. Yang H, Yuan H, Zhao X, Xun M, Guo S, Wang N, Liu B, Wang H. 2022. Cytoplasmic domain and enzymatic activity of ACE2 are not required for PI4KB dependent endocytosis entry of SARS-CoV-2 into host cells. Virologica Sinica 37:380–389.

50. Tada T, Fan C, Chen JS, Kaur R, Stapleford KA, Gristick H, Dcosta BM, Wilen CB, Nimigean CM, Landau NR. 2020. An ACE2 Microbody Containing a Single Immunoglobulin Fc Domain Is a Potent Inhibitor of SARS-CoV-2. Cell Rep 33:108528.

51. Hofmann H, Simmons G, Rennekamp AJ, Chaipan C, Gramberg T, Heck E, Geier M, Wegele A, Marzi A, Bates P, Pöhlmann S. 2006. Highly conserved regions within the spike proteins of human coronaviruses 229E and NL63 determine recognition of their respective cellular receptors. J Virol 80:8639–52.

52. Liu H, Naismith JH. 2008. An efficient one-step site-directed deletion, insertion, single and multiple-site plasmid mutagenesis protocol. BMC Biotechnology 8:91.

53. Liu S, Selvaraj P, Lien CZ, Nunez IA, Wu WW, Chou C-K, Wang TT, Subbarao K. 2021. The PRRA Insert at the S1/S2 Site Modulates Cellular Tropism of SARS-CoV-2 and ACE2 Usage by the Closely Related Bat RaTG13. Journal of Virology 95:e01751–20.

54. Paoloni-Giacobino A, Chen H, Peitsch MC, Rossier C, Antonarakis SE. 1997. Cloning of the TMPRSS2 gene, which encodes a novel serine protease with transmembrane, LDLRA, and SRCR domains and maps to 21q22.3. Genomics 44:309–20.

55. Naldini L, Blomer U, Gallay P, Ory D, Mulligan R, Gage FH, Verma IM, Trono D. 1996. In vivo gene delivery and stable transduction of nondividing cells by a lentiviral vector. Science 272:263–7.

56. Zufferey R, Nagy D, Mandel RJ, Naldini L, Trono D. 1997. Multiply attenuated lentiviral vector achieves efficient gene delivery in vivo. Nat Biotechnol 15:871–5.

57. Lee JM, Huddleston J, Doud MB, Hooper KA, Wu NC, Bedford T, Bloom JD. 2018. Deep mutational scanning of hemagglutinin helps predict evolutionary fates of human H3N2 influenza variants. Proc Natl Acad Sci U S A 115:E8276–e8285.

58. Wang W, Xie H, Ye Z, Vassell R, Weiss CD. 2010. Characterization of lentiviral pseudotypes with influenza H5N1 hemagglutinin and their performance in neutralization assays. J Virol Methods 165:305–10.

59. O’Doherty U, Swiggard WJ, Malim MH. 2000. Human immunodeficiency virus type 1 spinoculation enhances infection through virus binding. J Virol 74:10074–80.

60. Lin HX, Feng Y, Tu X, Zhao X, Hsieh CH, Griffin L, Junop M, Zhang C. 2011. Characterization of the spike protein of human coronavirus NL63 in receptor binding and pseudotype virus entry. Virus Res 160:283–93.

